# Force loading on molecular clutches governs the stability of cell lamellipodia

**DOI:** 10.1101/2025.10.10.679903

**Authors:** Ruihao Xue, Lezi Kang, Yonggang Chen, Haoxiang Yang, Hongyuan Jiang, Ze Gong

## Abstract

Cells can utilize the lamellipodia, a thin actin-rich membrane protrusion, to probe the mechanical properties of microenvironments. During the mechanosensing process, the lamellipodium usually exhibits instability in dynamics, *i.e*., protrusion-retraction cycles. However, how mechanical instability arises in lamellipodia, along with the functional role of dynamic instability in mechanosensing, is poorly understood. Here, we develop a minimal mechanochemical model for lamellipodia dynamics that integrates membrane deformation, myosin contractility, and binding kinetics of adhesion molecules (molecular clutches). Through stochastic simulations and analytical mean-field analysis, we demonstrate that both force-loading rate and magnitude applied by myosin-driven retrograde flow mediate the clutch binding kinetics, governing lamellipodial stability and hence the mechanosensing. Specifically, a slow force loading rate allows the clutches to bind and traction to accumulate, while a high loading magnitude collapses the bound clutches, causing protrusion-retraction cycles (instability) in lamellipodia. Our model predicts that a stiffer substrate stabilizes the lamellipodia by increasing the force loading rate, consistence with previous experiments. Furthermore, our model predictions on the biphasic regulation effect of myosin perturbations have been quantitatively validated by experimental results. Overall, the theoretical framework highlights the force loading as the key mechanical input driving lamellipodial instability and cell mechanosensing, advancing our understanding of mechanotransduction’s role in cell behaviors.

**Significance:** Cell behavior can be highly dynamic, showing periodic protrusion and retraction at the leading edge, and this dynamic instability is key to migration, immune response, and cancer invasion. How such instability arises from mechanotransduction between intercellular components and extracellular matrices remains unclear. Here, through theoretical modeling, we identify the rate and magnitude of force loading on adhesion molecules (molecular clutches) as the key factors for controlling lamellipodial stability. We find that a slow loading rate enables more clutches to bind and traction to accumulate, while a high loading magnitude collapses the bound clutches, causing protrusion-retraction cycles in lamellipodia. This work reveals a physical mechanism of how cells sense mechanical cues and adapt their dynamics, offering insights into diverse biological processes.

## Introduction

Instability has been widely observed in biological systems, from tissue morphogenesis to subcellular dynamics. During embryo development, mechanical instabilities drive essential shape transformations such as cortical folding (1, 2) and intestinal villus formation (3), enabling tissues to acquire functional architectures. At the cell level, similar instability phenomena, such as cyclic protrusion behaviors in invadosomes (4, 5), actin waves, and cortical chemical patterning (6, 7), have been implicated in cell migration, immune responses, and cancer progression (8).

Cells are able to probe the mechanical properties of the extracellular matrix (ECM) and actively regulate cell behaviors (9–12). This cell mechanosensing capability requires the lamellipodium, a thin actin-rich membrane protrusion at the leading edge of spreading cells, where mechanical forces transmit from intracellular components (*e*.*g*., myosin, cytoskeleton) to the ECM via adhesion molecules. Studies have shown that the lamellipodial dynamics also exhibit instability, *i*.*e*., membrane ruffling and protrusion-retraction cycles (13, 14). Importantly, lamellipodia can actively vary their dynamic pattern in response to changes in extracellular substrate stiffness and fluid viscosity (15–17). Therefore, understanding how such mechanical instabilities arise is essential for uncovering the mechanisms of cellular mechanosensing.

The molecular clutch concept has been widely adopted to depict the stochastic adhesion bond kinetics between the actin cytoskeleton and the ECM (18–20). By combining the molecular clutch with the myosin motor contractility, computational studies established the motor-clutch model to understand cell mechanosensing of substrate stiffness and viscoelasticity (21–24). Recent studies have shown that forces loading on molecular clutch can influence cell spreading and direct mesenchymal stem cells differentiation by fabricating nanopatterned substrates with varied integrin ligand spacing (25, 26). Additionally, using DNA-based tension probes, researchers have measured the rate of force loading experienced by molecular clutches and identified the loading rate as essential for mechanotransduction (27–29). Despite these advances, a theoretical framework that can be used to interpret how the magnitude and rate of force loading shape the lamellipodial instability and mechanosensing remains lacking.

Here, we develop a minimal mechanochemical model based on the motor-clutch framework by integrating the myosin contractility, molecular clutch stochastic binding kinetics, membrane deformation, and force-mediated actin polymerization. Our model identifies the force-loading rate and magnitude on molecular clutches as the key factor that governs lamellipodial stability and hence mechanosensing. Specifically, a slow force-loading rate allows the clutches to bind and traction accumulate to support protrusion, while a high force-loading magnitude causes the catastrophic failure of bound clutches and induces retraction; their temporal interplay gives rise to protrusion-retraction cycles (instability) in lamellipodia. In contrast, when force loading is both fast and strong, clutch failure occurs quickly, leading to persistent retrograde actin flow and insufficient protrusion; weak force loading stabilizes adhesions and enables steady protrusion. Based on this, three distinct dynamical regimes: stick-slip, sliding, and stick-locked dynamics, with the analytical critical force loading, are identified. Notably, our model predicts transitions between these distinct lamellipodial dynamics in responses to different ECM stiffness and pharmacological or genetic modulation of myosin activity, consistent with experimental observations. Our theoretical framework reveals the mechanism of how cells utilize the force loading to integrate intra- and extra-cellular mechanical cues, regulating their lamellipodial stability and adapting to their mechanical environment.

## Model and Results

### Kinetics of molecular clutch and cell spreading

During cell spreading, globular actin (G-actin) rapidly polymerizes to form actin filaments (F-actin) within the lamellipodium, generating polymerization forces that push and bend the cell membrane (Fig. 1A-B). To facilitate the model establishment, we assume that the cell spreads in a circular shape, expanding from an initial radius *r*_0_ to a spreading radius *r*, and the lamellipodium has a thickness of 2*h*. According to the motor-clutch hypothesis (21, 30, 31), the membrane resistance *f*_mem_ and myosin II (MII) contraction *f*_myo_ collectively drive the retrograde flow of F-actin toward the cell center (see Fig. 1C). Concurrently, molecular clutches transmit these forces from F-actin bundles to the ECM by stochastically engaging and disengaging with the actin cytoskeleton, which allows cells to dynamically probe the underlying ECMs.

**Fig. 1.**
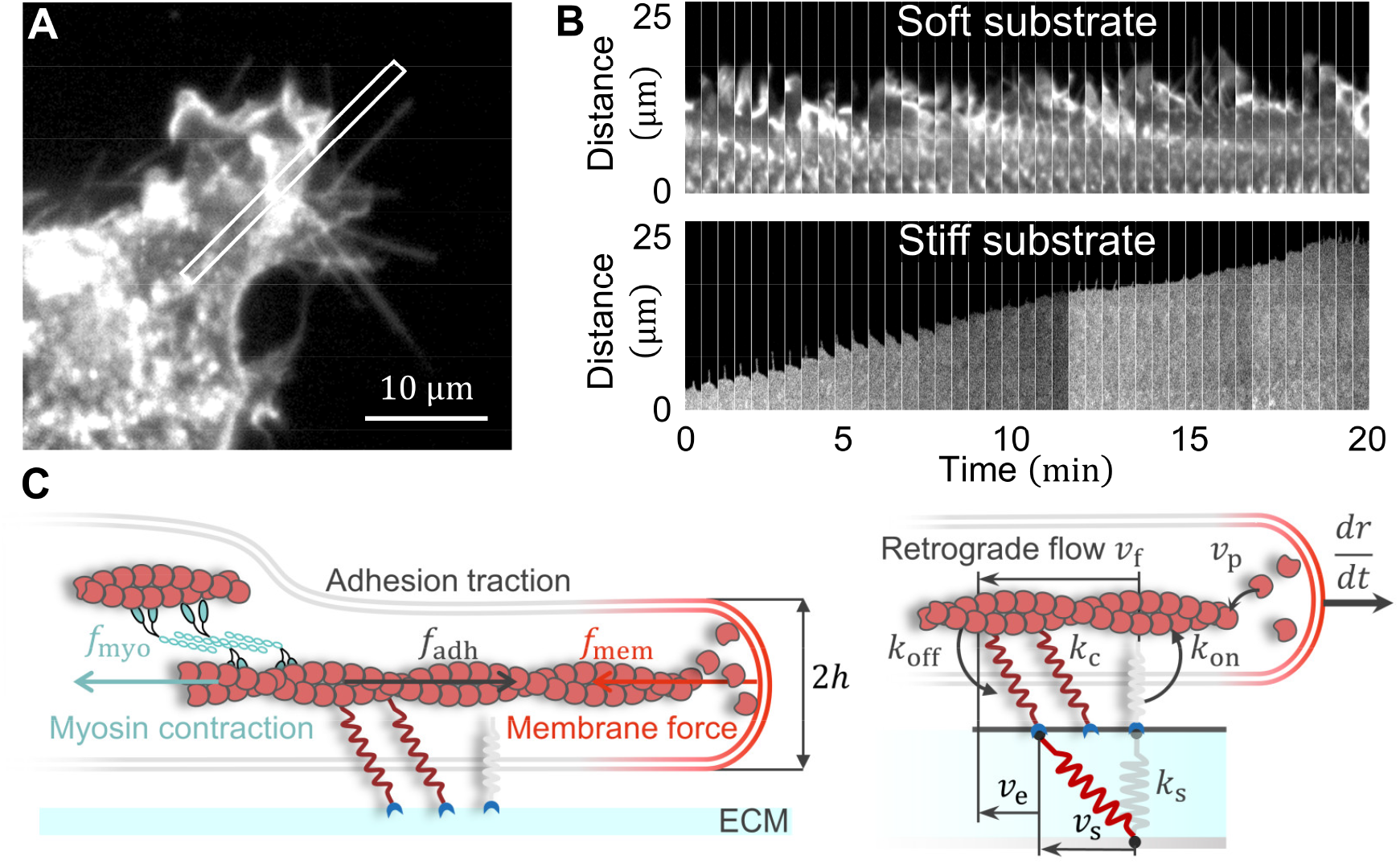
Sketch of the model describing lamellipodia dynamics. (**A**) Fluorescence image of a fibroblast expressing a GFP membrane marker, cultured on a compliant substrate. Scale bar: 10 μm. (**B**) Time-lapse montage of the boxed region in panel (**A**), showing cyclic protrusion– retraction on soft substrates (2.1 kPa) and persistent forward protrusion on stiff substrates (48 kPa). (**A**-**B**) Experimental images adapted from Ref. (16). (**C**) Schematic of the conventional motor-clutch model. The cell spreads radially with a lamellipodium of thickness 2*h* . Actin polymerization at a speed *v*_p_ pushes the membrane forward, while membrane resistance *f*_mem_ and MII contraction *f*_myo_drive F-actin retrograde flow at speed *v*_f_. Molecular clutches (stiffness *k*_c_) stochastically link F-actin (on-rate *k*_on_, off-rate *k*_off_) to the ECM, transmitting traction forces *f*_adh_ to deform the substrate (stiffness *k*_s_). The net cell spreading speed is *dr*/*dt* = *v*_p_ − *v*_f_. The geometric relation between the substrate deformation speed *v*_s_, clutch extension speed *v*_e_, and retrograde flow speed *v*_f_, satisfying *v*_s_+ *v*_e_ = *v*_f_.

#### Adhesion dynamics

We suppose that an adhesion consists of *N*_*c*_ assembled molecular clutches, each modeled as linear springs with stiffness *k*_*c*_. When engaged, the upper ends of the clutches attach to actin filaments and move centripetally at a retrograde flow speed *v*_f_, while their lower ends stretch the elastic substrate (with stiffness *k*_s_). Following Bell’s model (32), the clutch force *f*_c,*i*_ increases, leading to an exponential rise in the dissociation rates 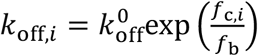. Here *i* = 1, ⋯, *N*_c_ indicating the *i* th clutch, 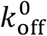 is the initial dissociation rate and *f*_b_ is a characteristic force. Once a clutch disengages, the clutch returns to a relaxed state (with zero extension) such that its upper end follows substrate displacement *u*_s_; this displacement is given by 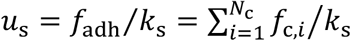 considering the balance between the total adhesion force 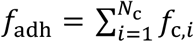 and the substrate elastic force. Following disengagement, it rebinds to F-actin at a constant association rate *k*_on_, undergoing stochastic engagement-disengagement cycles.

The adhesion traction force *f*_adh_ generated by engaged clutches resists the retrograde flow of F-actin, while the cell membrane at the lamellipodium provides a resistance force *f*_mem_ that opposes cell spreading (Fig. 1C). Based on force equilibrium on F-actin, the myosin contraction force *f*_myo_ is given by *f*_myo_ = *f*_adh_ − *f*_mem_. By applying the linearized Hill’s relation (21) of 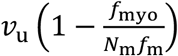, the retrograde flow speed *v*_f_ is obtained as

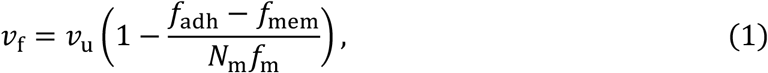

where *N*_m_ is the total number of myosin motors in the F-actin bundles, *f*_m_ is the characteristic stalling force per myosin, and *v*_u_ represents the unloaded retrograde flow speed. When adhesion molecules completely “clutch” the F-actin bundle (*v*_f_ = 0), myosin motors exert their maximal contraction force of *N*_m_*f*_m_.

#### Membrane deformation

To calculate the membrane force *f*_mem_, we employ a free energy framework (33, 34) in which the cytoskeleton is modeled as a homogeneous, isotropic medium composed of *N* F-actins that collectively support the membrane. The total work performed by *N* F-actins within lamellipodium is *G*_load_ = −*Nf*_actin_(*r* − *r*_0_), where *f*_actin_ is the force sustained by each filament. Meanwhile, the membrane is subject to stretching and bending due to actin polymerization. To model the free energy contributed by stretching and bending, we considered the membrane area change Δ*A*, and the free energy associated with membrane stretching is *G*_stretch_ = *σ*Δ*A* with *σ* representing the surface tension; The membrane bending energy can be expressed in the Helfrich–Canham–Evans’s form (35–37): *G*_bend_ = ∫ 2*κH*^2^*dA*, where *κ* represents bending stiffness and *H* is the mean curvature of membrane. The total free energy can be written as *G*_tot_ = *G*_load_ + *G*_stretch_ + *G*_bend_. By minimizing the total free energy with respect to radius *r*, we obtain and membrane resistance force acting on each filament as (refer to SI Note 1)

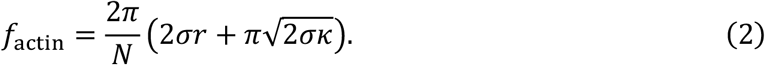

In addition, by minimizing *G*_tot_ with respect to *h*, we can obtain the half lamellipodium thickness 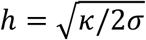. Under physiological conditions of *σ* = 0.02 pN/nm and *κ* = 80 p*N* · nm (37, 38), we find 2*h* ≈ 100 nm, which aligns with previous theoretical derivation and experimental observations (39–41). Note that, since membrane bending and stretching at filament tips contribute negligibly (*i*.*e*., 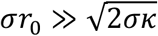), the force supported by each filament can be further approximated as *f*_actin_ ≈ 4*πσr*/*N*. Considering each adhesion connects *M* actin filaments that constitute the actin regulatory layer, the total membrane force transmitted through clutches becomes *f*_mem_ = *Mf*_actin_.

#### Cell spreading dynamics and non-dimensionalization

At the leading edge, actin polymerizes at a speed *v*_p_, yielding a cell spreading speed *dr*/*dt* = *v*_p_ − *v*_f_. Following the Brownian ratchet (BR) model (42), the actin polymerization is regulated by thermal fluctuations of the cell membrane under mechanical load. Considering that the local membrane sustains a local force *f*_actin_ from each actin filament, this mechanical load exponentially decreases the likelihood that the membrane thermal fluctuations create a sufficient gap (half-monomer size *δ* ) for a monomer to insert. Based on the Boltzmann distribution, the polymerization speed at each filament tip is 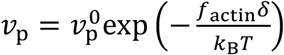. Here, 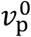 corresponds to the load-free polymerization speed. Note that, following previous studies (21, 22, 43, 44), *v*_u_ is typically set equal to 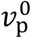 to prevent spontaneous cell spreading or retraction in the absence of adhesion traction. Combining Eqs (1) and (2), we derive the spreading speed as:

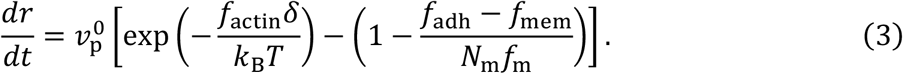

To further reduce the model complexity, we non-dimensionalize our model by using the time scale 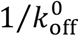, force scale *N*_c_*f*_b_, stiffness scale *N*_c_*k*_c_, and length scale *h*, such that we have 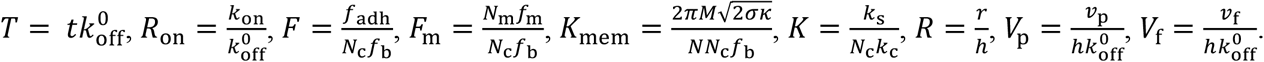.

Undering this scaling, the spreading equation becomes:

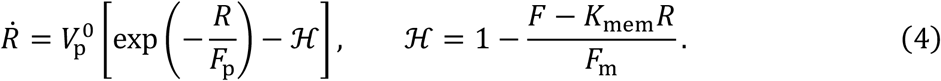

Here, the overdot denotes *d*/*dT*, and 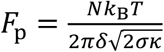is the characteristic polymerization force. For a detailed description of the model implementation and parameter settings, refer to SI Notes 1 and 2. The overall computational workflow is presented in Fig. S1.

### Cell mechanosensing requires the membrane tension-mediated Brownian ratchet at lamellipodia tips

By solving the BR-integrated motor-clutch model with the Gillespie algorithm (45), we investigate how cells respond to substrate stiffness by examining their spreading processes across a broad range of substrate stiffnesses. At low to intermediate stiffness ( *K* = 10^−4^ to 10^−2^ ), enhanced adhesion traction promotes cell spreading, leading to an increase in cell radius *R* (Fig. 2A). This expanding lamellipodium increases membrane resistance, which suppresses G-actin polymerization speed *V*_p_ via BR mechanism. The reduced polymerization further slows down F-actin retrograde flow *V*_f_, which in turn increases myosin contractility *F*_myo_= *F*_m_(1 − *ℌ*) via Hill’s relation (Fig. 2B).

**Fig. 2.**
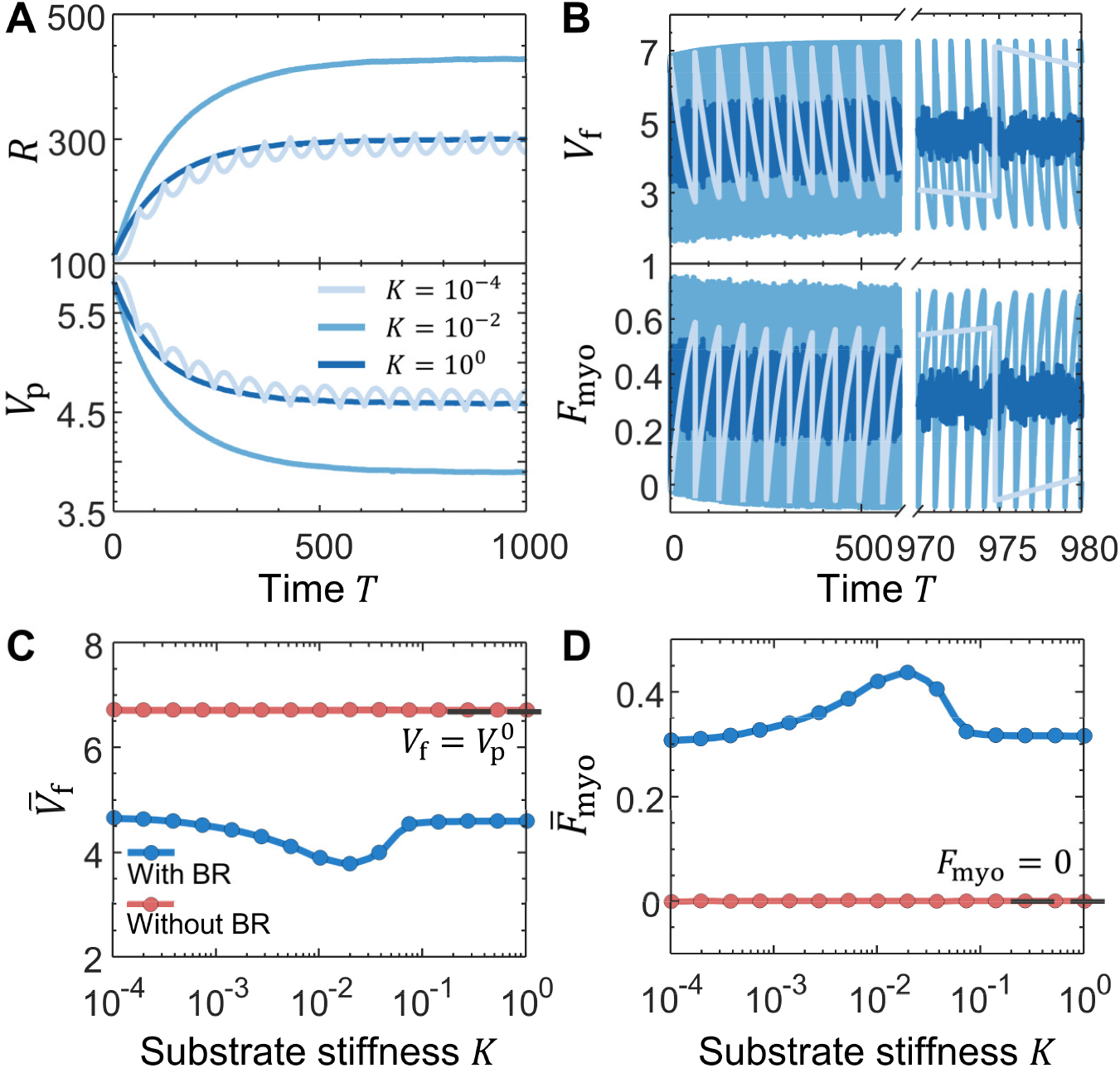
Substrate stiffness and myosin contractility regulate cell spreading dynamics. (**A-B**) Temporal profiles of key parameters in the BR-integrated motor-clutch model under increasing substrate stiffness *K*: (**A**) cell spreading radius *R* and membrane-regulated actin polymerization speed *V*_p_ ; (**B**) retrograde flow *V*_f_ and myosin contractility force *F*_myo_. (**C**-**D**) The motor-clutch models with BR (blue curves) and without the BR model (red curves) predict (**C**) retrograde flow speed 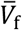, and (**D**) myosin force 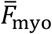, as a function of *K*. A bar over a variable denotes a time-averaged value. Parameters: (**A-D**) *F*_m_ = 1.

To clarify the role of the membrane-regulated polymerization, we compared cell behaviors from the conventional motor-clutch model (22, 43, 44), in which actin polymerizes at a constant rate 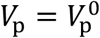. In this model, both the time-averaged retrograde flow speed 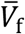 and myosin force 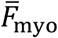 remain nearly constant across a broad range of substrate stiffness *K* (Fig. 2C-D, red curves). This arises because, in steady state, retrograde flow matches the fixed polymerization speed 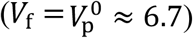, resulting in a uniform retrograde flow speed. Consequently, myosin generates the same level of contraction to maintain this uniform flow, regardless of the substrate stiffness *K*. While this simplification provides insight into basic clutch dynamics, it fails to reproduce the experimentally observed variations in retrograde flow and myosin activity across diverse mechanical properties of ECMs (10, 25, 26, 46). In contrast, the BR-integrated model captures mechanical feedback between adhesion and membrane tension, yielding distinct yet closely related biphasic responses in both 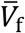 and 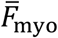 via BR (Fig. 2C-D, blue curves). Both quantities peak at intermediate stiffness values, aligning with previous experimental observations, highlighting the importance of tension-mediated regulated polymerization in cell mechanosensing (10, 21, 47).

In addition, our BR-integrated model predicts two distinct spreading dynamics regulated by substrate stiffness. On a soft substrate (Fig. 2A), molecular clutches are unstable. They repeatedly load and release the substrate, resulting in cyclic protrusions-retractions in the cell radius *R*. In contrast, on a stiff substrate (Fig. 2A), as molecular clutches sustain higher tensions, they stochastically bind and rapidly detach, generating a stable traction to support the cytoskeleton, thereby allowing the cell radius to vary smoothly over time. These stiffness-dependent dynamics agree with both classic motor-clutch model predictions and experimental observations (Fig. 1B) (15, 16, 21). To address the underlying physical mechanisms governing the dynamics, we next study how the stochastic binding and breaking kinetics of molecular clutches in our BR-integrated model are affected by substrate stiffness.

### Mean-field approximation for the BR-integrated motor-clutch model

We first applied the mean-field theory to approximate the BR-integrated motor-clutch model. Inspired by previous derivations (48–50), we begin by approximating the adhesion traction force through force balance with the substrate: *f*_adh_ = *k*_s_*u*_s_. Based on the geometrical relation illustrated in Fig. 1C, the combined extension of the substrate spring *u*_s_ and bound clutches *u*_e_ equals the displacement induced by actin retrograde flow *u*_*f*_, i.e., *u*_s_ + *u*_e_ = *u*_f_ . Thus, we reformulate the traction as *f*_adh_ = *k*_s_(*u*_f_− *u*_e_), yielding the traction dynamics: *df*_adh_/*dt* = *k*_s_(*v*_f_− *v*_e_), where *v*_f_ is defined by Hill’s relation (Eq. 1), and *v*_e_ represents clutch stretching speed. Assuming *N*_b_ engaged molecule clutches sustaining a uniform traction *f*_*c*_ = *k*_*c*_*u*_e_, we consider that they get stretched at a speed *v*_e_ over their lifetime 1/*k*_off_. This gives the average extension *u*_e_ = *v* /*k*_off_ such that we can obtain 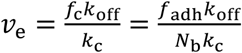 . For slip bonds, the mean-field approximation for the off-rate is written as 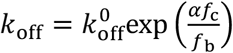, where the correction coefficient *α* = 1.35 is introduced to compensate for the underestimation of the off-rate in the mean-field approximation. Following our previous non-dimensionalizing, we define dimensionless clutch force 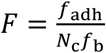, the bound probability 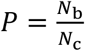, and the cell radius 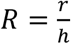. This leads to a reduced system of three coupled ODEs:

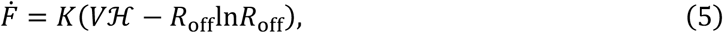

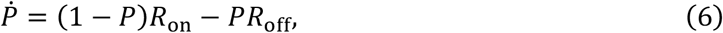

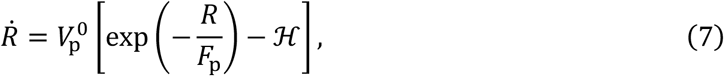

where we have the dimensionless speed 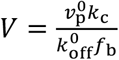, rescaled Hill’s relation 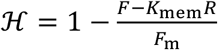, and off-rate 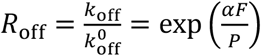 . Eq. (6) corresponds to the master equation for the bound probability, and Eq. (7) governing spreading dynamics was previously derived (see Eq. 4) but is included here for completeness.

Considering 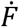governs the evolution of traction, we now interpret the two terms on the right-hand side of Eq. (5). The first term *KVℌ* represents the load from actin retrograde flow, while the second term *KR*_off_ln*R*_off_ accounts for traction loss due to clutch disengagement. We found that traction fails catastrophically when this loss term overtakes the actin-driven load term, which occurs when *F*/*P* > *W*(*Vℌ*)/*α*. Here, *W* is the Lambert *W* function defined by *x* = W*e*^*W*^, and *F*/*P* corresponds to the force-loading magnitude acting on each engaged clutch. For analytical clarity, we define the rupture threshold *F*\*: = W(*Vℌ*)/*α* ≈ W(*V*)/*α*, which marks the onset of bond failures. Since the clutch binding timescale can be characterized by 1/*R*_on_, the loading force experienced by newly engaged clutches can be approximated by *KVℌ*/*R*_on_ . Comparing this to the *F*\* allows us to evaluate whether the actin-driven loading rate *KVℌ* is fast enough to trigger bond failures before sufficient traction is accumulated. We thus define a critical loading rate *L*\*: = *W*(*V*)*R*_on_/*α* ≈ 13.71. Depending on the relative magnitude between *KVℌ* and *L*\*, the adhesion dynamics show three distinct behaviors:

a) *KVℌ* ≪ *L*\*: Slip-bond unbinding is negligible, and traction accumulates gradually.
b) *KVℌ*∼*L*\*: Slip-bond kinetics become significant, giving rise to nonlinear behaviors.
c) *KVℌ* ≫ *L*\*: Slip-bond unbinding dominates immediately, preventing traction build-up and leading to F-actin sliding smoothly.

Altogether, this analysis demonstrates that the actin-driven loading rate *KVℌ* is the key variable governing clutch behavior and traction evolution.

### Mean-field approximation yields different cell dynamics regulated by substrate stiffness

Next, we utilized the critical loading rate *L*\* to examine the effect of substrate stiffness on system stability. By applying a small perturbation (*δF, δP, δR*) to the fixed point (*F*_0_, *P*_0_, *R*_0_), we can examine the eigenvalues *λ* of the Jacobian matrix and plot the *K*-*F*_m_ phase diagram. Based on the phase diagram, we can categorize the dynamics into three types of behaviors as below:

(I) On a soft substrate, the actin-driven loading rate satisfies *KVℌ* ≪ *L*\* (Fig. 3C(*I*)). In this regime, slip-bond unbinding is initially negligible, allowing molecular clutches to maintain a high binding probability. Thus, the clutch force *F* increases quasi-linearly at a low loading rate 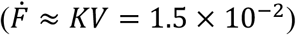, and each bound clutch shares a relatively low force (given by *F*/*P*). When *F*/*P* exceeds threshold *F*\* (dashed grey line in Fig. 3C), the failure of one certain engaged clutch triggers a cascade of bond failures, leading to a sudden drop in the adhesion force. As a result, this instability drives the cell radius to undergo repeated cycles of spreading and retraction (Fig. 2A).
(II) As substrate stiffness increases, the actin-driven loading rate *KVℌ* ≈ 5 approaches our critical value *L*\* (Fig. 3C(*II*)). In this intermediate regime, the nonlinear kinetics of slip-bonds take effect, driving cell dynamics into sustained oscillations. Specifically, the system undergoes a supercritical Hopf bifurcation at a critical stiffness *K*\*, where the real parts of the conjugate pair cross zero (51, 52) (Fig. 3B). This bifurcation raises a stable limit cycle that confines oscillatory behaviors (Fig. S2). The increased traction loading rate also shortens the oscillation period. These unstable, oscillatory behaviors (I, II) are known as stick-slip (or load-and-fail) dynamics (21, 49, 50, 53). Threshold *F*\* delineates the transition between adhesion “stick” states and “slip” failures.
(III) On a stiff substrate, the high stiffness induces a rapid clutch loading rate (*KVℌ* ≫ *L*\*). Once the molecular clutches engage, the off-rate increases immediately, preventing sustained engagement and the accumulation of substrate deformation (Fig. 3D(*III*) and 3B). As a result, clutches bind F-actin and release it rapidly, allowing F-actin to slide smoothly, which is also referred to as sliding dynamics or frictional slippage (21, 49, 53).

**Fig. 3.**
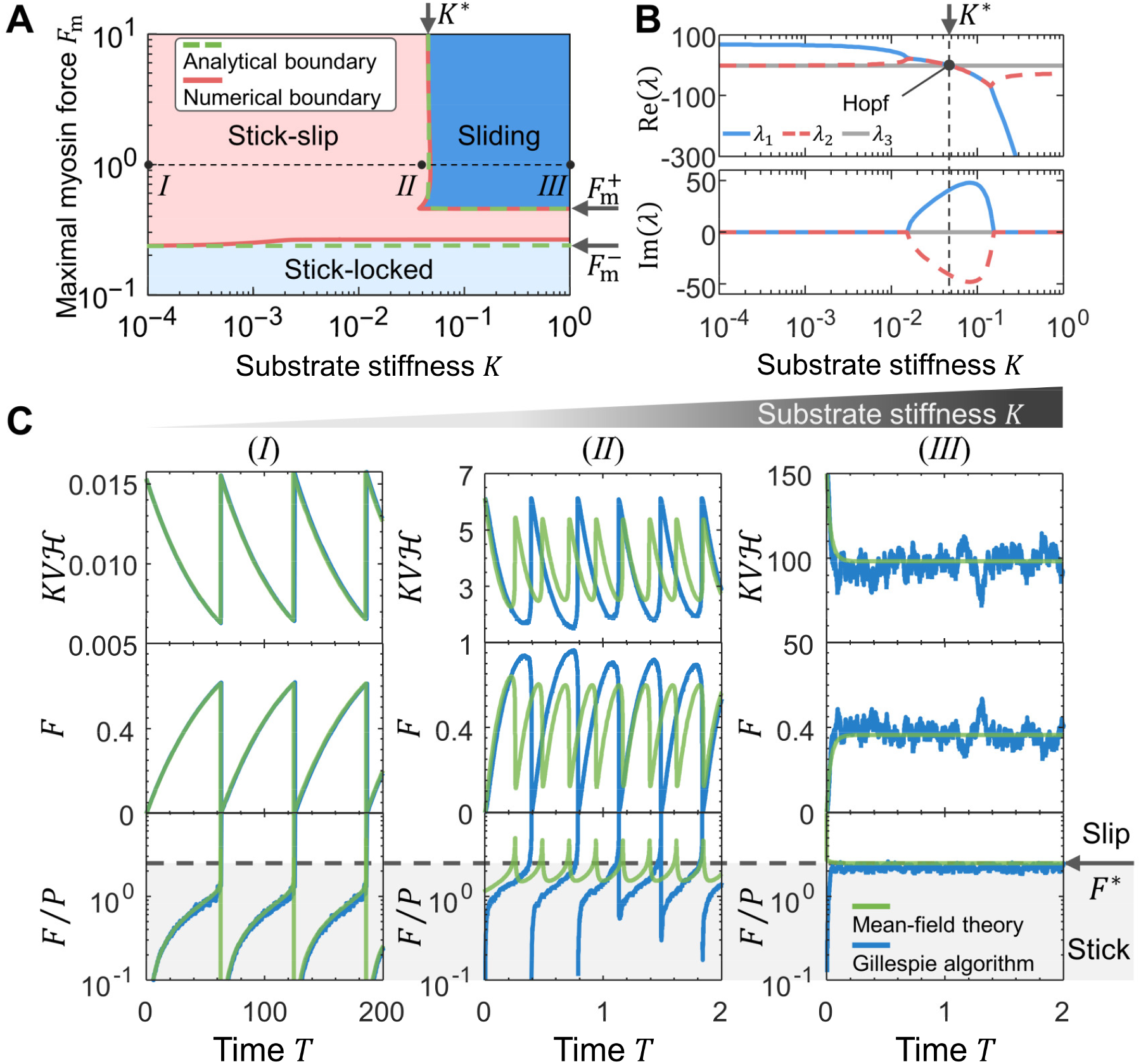
Stiffer substrate stabilizes adhesion by increasing the force-loading rate. *KV****ℌ***. (**A**) Phase diagram showing three distinct dynamical regions in the *K*-*F*_m_ parameter space: Sliding dynamic (blue): the traction force and cell radius reach steady-state values immediately, while F-actin rapidly slides toward the cell center. Stick-slip dynamic (pink): the system exhibits oscillatory behaviors and protrusion–retraction cycles. Stick-locked dynamic (light blue): the cell forms stable adhesions, with F-actin stably stalled by molecular clutches and gradual cell spreading. The stability boundary (red lines) is computed via numerical linear stability analysis. Analytical stability boundaries 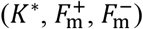 are derived from Eqs. (8)-(9). Black dots labeled with Roman numerals mark the parameter sets (*K, F*_m_): (*I*) (10^−4^, 10^0^); (*II*) (4 × 10^−2^, 10^0^); (*III*) (10^0^, 10^0^). (**B**) Real part Re(*λ*) and imaginary part Im(*λ*) of eigenvalues are plotted as functions of substrate stiffness. The black dot marks the critical stiffness *K*\* at the Hopf bifurcation. Parameter: *F*_m_ = 1. (**C**) Temporal dynamics of actin-driven loading rate *KVℌ*, clutch force *F*, and the sustained force per engaged clutch *F*/*P* comparing the mean-field results (green curves) and stochastic simulations (blue curves). Rupture threshold *F*\* ≈ 2.74 (gray dashed line) delineates the stick (sub-threshold) and slip (super-threshold) dynamics. Roman numerals correspond to parameter sets in panel (**A**).

To analytically estimate the critical stiffness *K*\*, we first note from numerical observations that the rescaled Hill term satisfies *KVℌ* ≈ *KV* across a wide range of *K* (Fig. 3C). Based on this observation, we approximate the system (Eqs. 5-7) by treating *ℌ* = 1, leading to a two-dimensional system: 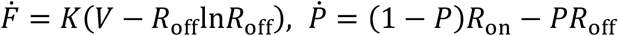. Performing linear stability analysis on this reduced system yields the critical stiffness (see SI Note 3 for derivation):

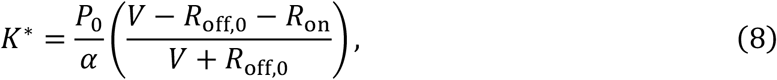

with *R*_off,0_ = *e*^*W*(*V*)^ and *P*_0_ = *R*_on_/(*R*_on_ + *R*_off,0_), which gives *K*\* ≈ 4.46 × 10^−2^, in excellent agreement with the numerically determined boundary (Fig. 3A).

### Myosin contraction regulates lamellipodium dynamics via force loading on molecular clutches

Considering that myosin contractility also affects the force loading, we next examine how myosin contractility impacts lamellipodial stability. As contractility decreases, the lamellipodium displays a series of dynamics: initially exhibiting insufficient spreading, then undergoing periodic protrusion-retraction cycles, and finally achieving steady protrusion (Fig. 4A). These dynamic patterns of lamellipodia are controlled by the actin-driven loading rate *KVℌ* (compared to the critical loading rate *L*\*), which determines whether molecular clutches can build up traction before detachment. High contractility generates fast retrograde flow, resulting in a rapid loading rate *KVℌ* ≫ *L*\* (Fig. 4B, left). In this case, clutches detach almost immediately after engaging, preventing traction buildup and resulting in continuous F-actin sliding. At intermediate contractility, molecular clutches sufficiently stall retrograde flow, allowing *KVℌ* to periodically drop below *L*\* (Fig. 4B, middle). When *KVℌ* ≲ *L*\*, molecular clutches bind to F-actin and collectively build up substantial traction to support membrane expansion (Fig. 4A). As the cell spreads, the clutch force gradually accumulates until the force per bound clutch *F*/*P* exceeds the slip-bond threshold *F*\*, triggering catastrophic adhesion failures (Fig. S5). This abrupt unbinding causes the membrane and myosins to propel actin filaments back toward the cell center, leading to a rapid lamellipodial retraction. When myosin contraction is sufficiently low, the actin-driven loading rate remains well below the critical value throughout the spreading process, *i*.*e*., *KVℌ* ≪ *L*\* (Fig. 4B, right). In this regime, the total internal force *F*_myo_+ *K*_mem_*R* becomes too small to rupture adhesion bonds, resulting in the clutch locking with F-actin. We refer to this newly identified regime as the stick-locked dynamic (Fig. 3A).

**Fig. 4.**
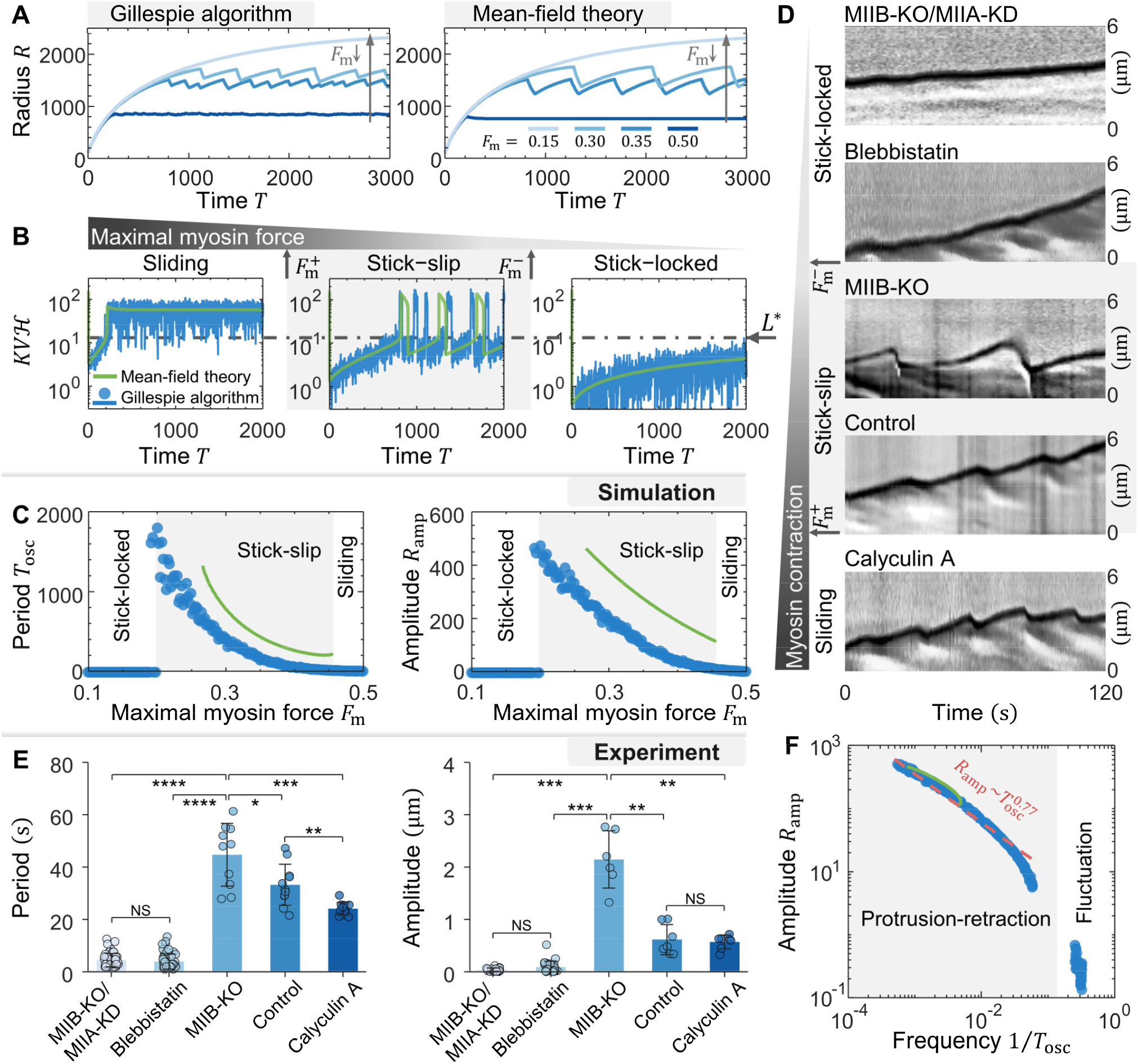
Myosin contractility tunes lamellipodial dynamics via force loading on molecular clutches. (**A**) Temporal evolution of cell radius *R* under increasing myosin contractility *F*_m_, showing three distinct stages of lamellipodial dynamics regulated by *F*_m_ . Left: stochastic simulation; right: mean-field approximation. (**B**) Actin-driven loading rate *KVℌ* under varying *F*_m_ as a function of time. The dashed-dotted line is the critical loading rate *L*\*. 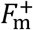 and 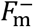 denote the upper and lower bounds of the stick-slip dynamic region, respectively. (**C**) Protrusion period *T*_osc_ (left) and amplitude *R*_amp_ (right) plotted as functions of *F*_m_ . (**D**) Kymographs show that increasing myosin activity via Calyculin A treatment shortens the oscillation period compared to controls, whereas MIIB-KO cells exhibit longer periods and larger protrusion amplitudes. MIIB-KO/MIIA-KD and blebbistatin-treated cells fail to generate periodic contractions. (**E**) Bar plots quantifying lamellipodial oscillation period (left) and amplitude (right) extracted from genetic or pharmacological modulation of myosin activity. Sample sizes of oscillation measurements (left to right): (left panel) 51, 56, 10, 11, and 11. (right panel) 26, 29, 6, 7, and 7. All error bars represent mean ± standard deviations (SD). Statistical analysis was performed using a two-sided Mann-Whitney *U*-test: ∗ *P* < 0.05,∗∗ *P* < 0.01,∗∗∗ *P* < 0.001,∗∗∗∗ *P* < 0.0001; NS: not significant. All experimental data in (**D-E**) are reproduced from Ref. (54). (**F**) Plotting *R*_amp_ versus 1/*T*_osc_ obtained from the BR-integrated model reveals an inverse power-law relationship. (**B**,**C**,**F**) Green curves show mean-field results; blue curves and scatter points represent stochastic simulations. Parameters: (**A-C**,**F**) *K* = 1; (**B**) left to right: *F*_m_ = 0.50, 0.35, 0.15.

To analytically estimate the lower critical myosin contractility 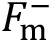, we first identify the maximal sustainable traction of the clutch 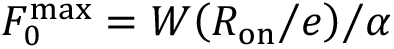, obtained from a geometric analysis of nullclines (Fig. S3). We next utilize the force balance on F-actin at the fixed point, where the clutch force satisfies: *F*_0_ = *F*_m_(1 − ℋ_0_) − *K*_mem_*F*_p_lnℋ_0_ (rearranged from Eqs. 5 and 7), and solve for *F*_m_ as a function of ℋ_0_:

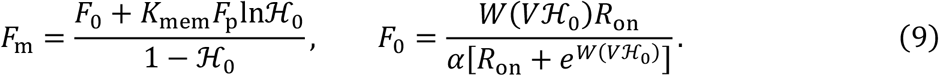

Here, *ℌ*_0_ denotes the Hill’s function at a stationary state. Under the critical condition 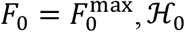can be derived as 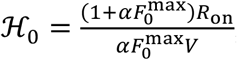, which yields the lower critical contractility 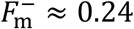. For the upper critical myosin contractility 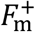, we analyze the bifurcation condition where the two nullclines become tangent at the fixed point (Fig. S4), marking the onset of stick-slip oscillations. Under this geometric condition, the Hill’s term satisfies *ℌ*_0_ ≈ 0.32, yielding the upper threshold 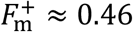 based on Eq. (9). Together, these analytical boundaries 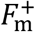 and 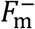 closely match the numerical transitions shown in Fig. 3A (Refer to SI Note 3 for more details).

### Experiments validate model-predicted biphasic regulation of lamellipodial dynamics to myosin contractility

To quantify our simulations, we extracted the oscillation amplitude *R*_amp_ and period *T*_osc_ of protrusion-retraction cycles, and we plot them as functions of myosin contractility *F*_m_ (Fig. 4C). Both quantities exhibit biphasic dependence on *F*_m_ . In the stick-locked and sliding regimes, molecular clutches stochastically engage and disengage with F-actin, causing small-amplitude, short-period fluctuations at the leading edge. Within the stick-slip regime, both *R*_amp_ and *T*_osc_ decrease progressively with increasing *F*_m_. By plotting the amplitude *R*_amp_ with frequency 1/*T*_osc_, we found an inverse power-law relationship between protrusion amplitude and frequency by both mean-field and stochastic approaches (Fig. 4F). Notably, the inverse amplitude-frequency relationship has also been reported in previous experiments (55).

To validate our model predictions, we closely examined a previous experimental study on lamellipodial retraction and adhesion formation, in which MII activity was selectively modulated through perturbations on fibroblast cells (54) (Fig. 4D). Surprisingly, their experiments also suggest the biphasic regulation of myosin on lamellipodial dynamics. By observing their kymograph data, we found that enhancing myosin activity with Calyculin A shortens the oscillation period, while MIIB knockout (MIIB-KO) leads to extended cycles with larger-amplitude; both are consistent with our model prediction. By additional knockdown of MIIA in MIIB-KO (MIIB-KO/MIIA-KD) cells, they found that cells displayed no visible oscillations in spreading (54). Since MIIA and MIIB together contribute to approximately 90% of total traction forces (56), the MIIB-KO/MIIA-KD cell fails to generate sufficient contractility to rupture adhesions and initiate retraction. Similarly, the blebbistatin treatment that inhibits MIIA and MIIB isoforms by suppressing ATPase activity (57, 58) also causes stable lamellipodia without oscillation. To quantitatively verify the biphasic response, we also extracted oscillation amplitudes and periods from the experimental kymographs for different myosin perturbation conditions (Fig. 4E). Interestingly, both amplitude and period increase under weak contractility, but are suppressed under the strong contractility case (Calyculin A treatment), collectively supporting the biphasic regulation of lamellipodial dynamics by myosin activity. Note that the measured fluctuations for MIIB-KO/MIIA-KD and blebbistatin-treated cells (stick-locked regime) are mainly driven by stochastic clutch kinetics and thermal noise at the cell membrane.

In addition to the oscillatory behaviors described above, we simulated the effects of pharmacological myosin inhibition on the overall spreading processes. In our model, reducing *F*_m_ leads to slower actin retrograde flow *V*_f_ (Fig. 5A), which enables adhesion traction to support the membrane resistance and promote steady-state spreading (Fig. 5B). Notably, our BR-integrated model quantitatively captured the experimentally observed increase in spreading area under different perturbations of myosin (56, 59, 60).

**Fig. 5.**
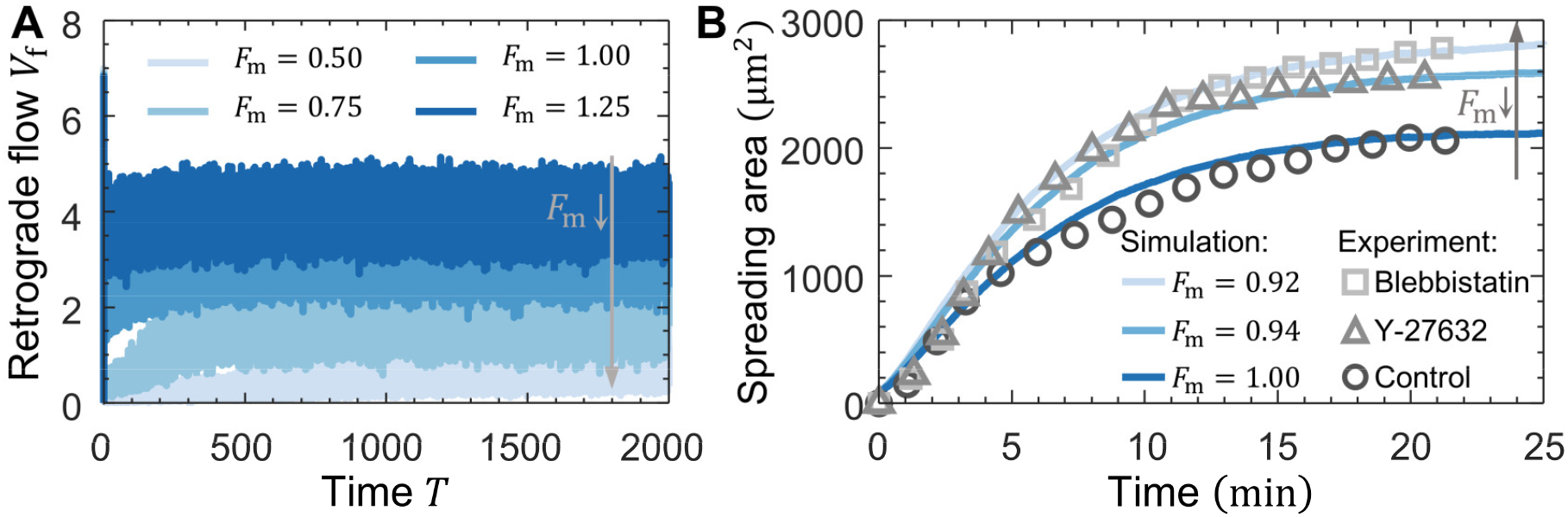
Lower myosin contractility increases spreading speed and area. (**A-B**) Temporal evolution of retrograde flow speed *V*_f_(**A**) and the spreading area *πr*^2^ (**B**) obtained from the BR-integrated model. Scatter plots in panel (**B**) represent the mean cell spreading area, extracted from Ref. (60). Grey arrows denote the shifts in the temporal trajectories on decreasing maximal myosin force *F*_m_. Parameters: *K* = 1, *R*_on_ = 8.4.

## Discussion and Conclusion

In this study, we developed a minimal mechanochemical model, named as BR-integrated motor-clutch model, that incorporates membrane tension-regulated actin polymerization and mechanical feedback between the cytoskeleton and the lamellipodial membrane. Our model reveals that increased substrate stiffness promotes cell spreading and elevates membrane resistance, which in turn suppresses G-actin polymerization via the BR mechanism, slows retrograde flow, and enhances myosin contractility via Hill’s relation. This mechanosensitive feedback enables the system to exhibit biphasic responses that align with experimental observations. Importantly, this framework overcomes a key limitation of conventional motor-clutch models, which fail to capture how retrograde flow and myosin activity respond to substrate stiffness at steady state (22, 43, 44). Furthermore, our model reproduces key trends observed under myosin inhibition, including enhanced spreading speed and increased cell size with reduced contractility.

Using linear stability analysis on the mean-field approximation of our model, we revealed three distinct dynamical regimes—sliding, stick-slip, and stick-locked. These distinct dynamics are governed by the two factors: the force-loading rate *KVℌ* that depicts how rapidly force is transmitted through engaged clutches, and the force-loading magnitude *F*/*P* that determines whether the transmitted force is sufficient to rupture adhesions and trigger retraction. In the sliding regime (Fig. 6A), a high loading rate ( *KVℌ* ≫ *L*\* ) and loading magnitude near the rupture threshold (*F*/*P* ≈ *F*\*) lead to continuous clutch failure and F-actin sliding toward the cellular center. This behavior emerges when both substrate stiffness and myosin contractility are high 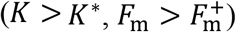. In the stick-slip regime (Fig. 6B), *KVℌ* ≲ *L*\*, allowing transient traction buildup. However, the internal contractile forces are still strong enough that the *F*/*P* eventually exceeds *F*\*, triggering periodic adhesion failure and oscillatory dynamics. This instability emerges when the substrate is soft (*K* < *K*\*) or myosin contractility is moderate 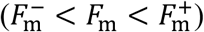. Finally, in the stick-locked regime (Fig. 6C), both *KVℌ* and *F*/*P* remain below their respective criteria, allowing stable clutch engagement and F-actin flow to be effectively stalled. This stable adhesion state arises when myosin contractility is sufficiently low 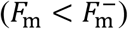. Note that a high loading rate without a corresponding force magnitude is incompatible, as rapid force-loading rate on the molecular clutch requires a sufficiently high myosin force to generate.

**Fig. 6.**
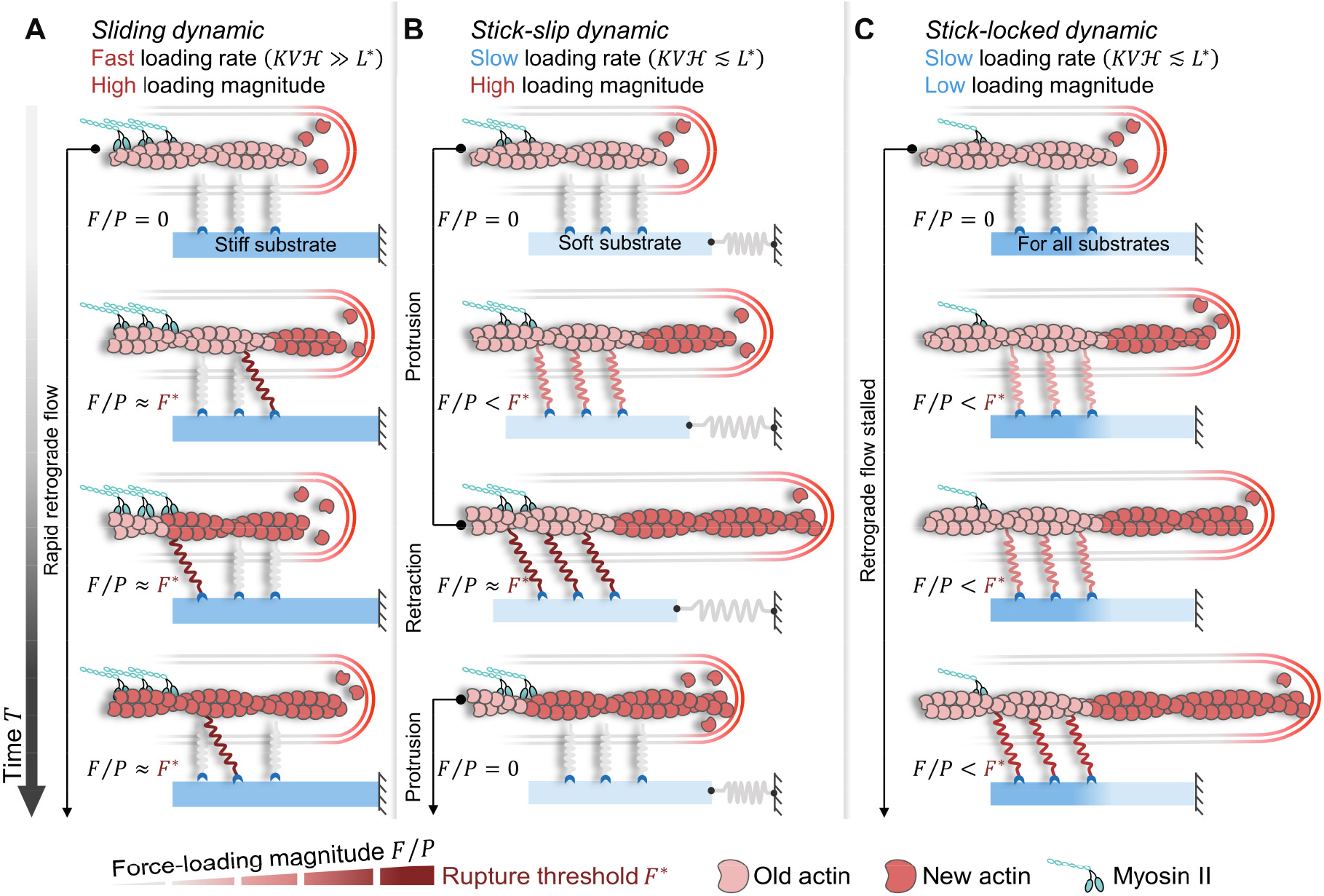
Schematic describes three types of lamellipodia dynamics. (**A**) Sliding dynamic: When both myosin contractility and substrate stiffness are high 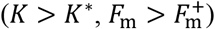, rapid force-loading rate ( *KVℌ* ≫ *L*\* ) leads to immediate clutch detachment. The force-loading magnitude is maintained near the failure threshold (*F*/*P* ≈ *F*\*), preventing traction buildup and resulting in fast retrograde flow. The lamellipodium exhibits small-amplitude, short-period fluctuations. (**B**) Stick-slip dynamic: For moderate contractility or soft substrates 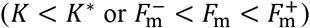, force-loading rate exhibits below the critical value *KVℌ* ≲ *L*\*, allowing traction to build before a sudden rupture occurs (*F*/*P* > *F*\* ), causing oscillatory protrusion-retraction cycles. (**C**) Stick-locked dynamic: When myosin contractility is low 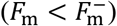, both the loading rate and force magnitude remain sub-threshold throughout spreading ( *KVℌ* ≲ *L*\*, *F*/*P* ≤ *F*\* ). Clutches remain stably locked, effectively stalling retrograde flow and enabling continuous protrusion.

Recent experiments have applied the DNA-based tension probe to measure force-loading rates on molecular clutches of 0.2 − 4 p*N*/*s* within focal adhesions on rigid substrates (27–29), which are markedly lower than those predicted by conventional motor-clutch simulations (typically 10^2^ − 10^4^ pN/s) (21, 22, 47, 61). Our model identified the critical force-loading rate *L*\* = W(*V*)*R*_on_/*α* and its loading rate scale 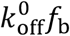, yielding a physical loading rate 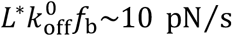. This indicates that traction can only be generated when the local force-loading rate remains below this kinetic threshold 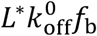. Accordingly, in focal adhesions where traction is stably established, the loading rate is constrained by clutch binding kinetics. In these regions, molecular clutches collectively stall retrograde flow, thereby keeping the force-loading rate below 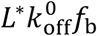 and maintaining adhesion architecture (clutch engagement). In contrast, Liu and colleagues (24) have attributed the low experimental values to the compliant elasticity of talin, with stiffness ranging from 0.05 to 1.6 pN/nm. Thus, our model provides an alternative explanation on the experimentally observed low loading rates without invoking complex elastic properties of adhesion proteins.

In summary, our theoretical model demonstrates that cells utilize force loading on molecular clutches to control their lamellipodial stability, and thus, their mechanosensing. The combined theoretical and experimental efforts show that variations in substrate stiffness and myosin contractility affect the force loading process to modulate the dynamic patterns of lamellipodia. While our framework captures key mechanosensitive responses and spreading instabilities, future extensions may address related cell behaviors by incorporating more detailed biochemical signaling modules (such as RhoA/ROCK pathways and integrin turnover) as well as nonlinear substrate mechanics, integrin ligand spatial distributions, and actin network bending. Our framework provides a fundamental understanding of force transmission and the related mechanical instabilities in spreading cells, offering mechanistic insights into instability behaviors broadly observed in immune, cancerous, and neuronal systems *in vivo*.

## Methods

### Numerical implementation

The mean-field equations form a stiff system of ODEs, for which we recommend using stiff ODE solvers, such as MATLAB’s **ode15s**. The initial conditions are as follows *F*(0) = 0, *P*(0) = 0, and *R*(0) = *r*_0_/*h*. For stochastic simulations, the state transitions of molecular clutches were simulated using the Gillespie algorithm (45), and time integration was performed using the forward Euler method. All stochastic simulations were performed for 5000 dimensionless time units to ensure the system reached a dynamic steady state. For each parameter setting, we first fit the spreading trajectory using an exponential equation to determine the characteristic time *T*_s_(Fig. S6). Time-averaged quantities (*e*.*g*.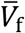 and 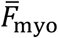) were then computed over the interval [5*T*_s_, 5000].

### Kymograph and montage generation

Kymographs and montage images were generated using *ImageJ* software (version 1.54p; https://imagej.net/ij/). To create kymographs, a straight line was drawn along the trajectory of the cell’s leading edge in time-lapse images, followed by the application of the “Reslice” function to produce spatiotemporal maps of membrane displacement. Montage images were assembled from the same time-lapse sequences using the “Make Montage” function, highlighting regions of interest that exhibited prominent morphological changes during the cell spreading process.

### Experimental data extraction

Oscillations of the leading edge (Fig. 4F-G) and cell spreading area (Fig. 5B) were extracted from experimental figures or kymographs using WebPlotDigitizer (version 5.2; https://automeris.io). Images were first imported into the website and calibrated by defining the X and Y axes according to known scale bars. To highlight the region of interest, the “Pen” tool was used to manually mark the signal trace as a mask. Data points were then extracted using the “Averaging Window” algorithm, which computes signal positions along the trace. The extracted data were exported as CSV files for subsequent analysis. Full stochastic simulation and quantification details can be found in SI Notes 2 and 4.

## Author contributions

R.X., H.J., and Z.G. designed research; R.X. and L.K. performed theoretical modeling, stability analysis, and numerical simulations; R.X., Y.C., and Z.G. analyzed data; R.X., Y.C., H.Y., H.J., and Z.G. wrote the paper.

## Data, materials, and software availability

All experimental data extracted from the published studies and our computational procedures are accessible at https://github.com/Tu2bo/Force-loading/tree/main. All other data are included in the main text and/or supporting information.

## Acknowledgments

This work was supported by the National Natural Science Foundation of China (Grants No. 12472323, No. 12202439, No. 12025207, and No. 11872357), the National Key Research and Development Program of China (2024YFF0509500), and the Fundamental Research Funds for the Central Universities. This work was partially carried out at the University of Science and Technology of China Center for Micro and Nanoscale Research and Fabrication.

